# COMPARATIVE PHENOMICS IN *Limnospira platensis* STRAINS REVEALS PHOTOBIOENERGETIC SIMILARITIES AND DIFFERENCES IN AMERICAN AND AFRICAN STRAINS

**DOI:** 10.64898/2026.09.25.754581

**Authors:** Mateo Zavala, Jose Armando Perez-Loera, Carli Peters, A Zavafer

## Abstract

This study addresses the functional diversity of wild *Limnospira platensis* strains from two continents. Wild strains retain regulatory and metabolic flexibility often lost during domestication, making them essential for exploring natural adaptation within the species’ evolutionary history. We investigated the comparative growth performance of five Limnospira strains under controlled temperature and light regimes to identify phenotypic differences attributable to single environmental variables. We then evaluated their biochemical and biophysical responses under naturally fluctuating light conditions, linking energy-use strategies to phylogenetic relationships. Our analyses reveal that while certain traits, such as fine-tuning of photosynthetic reactions, are shared among closely related strains, overall growth performance is highly variable—even within phylogenetically similar groups. For example, photosensitive strains (e.g., UTEX 3086) differ markedly from high-light tolerant strains (e.g., UTEX LB 2342) in their mechanisms of coping with light stress. Together, these comparative analyses highlight how environmental sensitivity and energy-use strategies vary across strains of the same species, reflecting broader patterns of diversification within Limnospira.

## 1. Introduction

The genus *Limnospira* (also known commonly as spirulina) has been actively studied due to their relevance in the food sector (Vieira et al. 2025). Conventional taxonomy considers this genus to be composed of four species: *L. platensis, L. maxima, L. indica and L. fusiformis* (Nowicka-Krawczyk et al. 2019; Roussel et al. 2023; Pinchart et al. 2024). The genus has a cosmopolitan distribution in the tropical and subtropical regions, being reported to occur in all continents (Roussel et al. 2023). While many strains have been isolated from salt lakes, some strains can be present in oceanic waters as well (Roussel et al. 2023).

Historically, Limnospira has been classified under the genera *Spirulina*, *Arthrospira*, and *Limnospira* (Sinetova et al. 2024); however, Limnospira is now considered the taxonomically correct genus (Nowicka-Krawczyk et al. 2019; Pinchart et al. 2024; Roussel et al. 2023). *Limnospira* has been hypothesized to have originated 8.2 MYA (based on Lifemap database, Jul/2026, see de Vienne (2016)) and is classified within the Oscillatoriales order (Ishida et al. 2001; Nowicka-Krawczyk et al. 2019) within the Sirenicapillariaceae family (Berthold et al. 2022). Spirulina strains display colonial filaments in the shape of a spiral, although this morphology of the trichomes seems to be heavily influenced by environmental conditions (Ma and Gao 2009), rather than a unique characteristic of Limnospira as a group (Nixdorf et al. 2003).

Most strains of *Limnospira* seem to prefer brackish to hypersaline waters, moderate to warm temperatures and waters rich in carbonate/bicarbonate, although, strains that occur naturally in fresh water have been reported (Roussel et al. 2023). The preference in water properties by species seems to be irrespective of their phylogenetic origin (Roussel et al. 2023). There are around 402 strains reported within the four species, the *Limnospira platensis* being the most diverse with at least 234 recognized strains (de Vienne 2016). However, different taxonomic analysis based on molecular data suggests that the separation between the four *Limnospira* species is artificial (Nowicka-Krawczyk et al. 2019; Roussel et al. 2023; Pinchart et al. 2024), and efforts to consider it a monospecific clade have been put forward to reclassify the whole clade as *L. platensis* (Roussel et al. 2023).

Despite the genus seeming to be genomically stable, there is a recognized phenotypic and metabolic diversity (Ogbonda et al. 2007; Roussel et al. 2023; Pinchart et al. 2024). The overwhelming majority of what we know about *L. platensis* growth comes from domesticated, commercially cultivated strains (Cheng et al. 2017; Benedetti et al. 2018; Xu et al. 2026). Thus, research tends to focus on multiple strains collectively, although the non-canonical strain PCC 7345 is one of the most studied (Rippka et al. 1979; Nelissen et al. 1994; Paramanya et al. 2024) and considered to be a taxonomic type for *L. platensis* by the NCBI (Taxonomy ID: 118562, Schoch et al. (2020)).

Beyond the biotechnological implications of the study of *L. platensis*, its photophysiology is remarkable. For example, at light harvesting level, *L. platensis*’s phycobilisomes have an unusually high level of C-phycocyanin (up to 18% of dry biomass) (Boussiba and Richmond 1980, Minkova et al. 2003). Saturation of growth is typically around 300–400 µmol photons m⁻² s⁻¹ (Singh and Singh 1997), O_2_ evolution maximum occurs close to 600 µmol photons m⁻² s⁻¹ (Vonshak et al. 1996) and sensitivity to photoinhibition is > 600 µmol photons m⁻² s⁻¹ (Jensen and Knutsen 1993), although this is highly strain-dependent (Vonshak et al. 1996; Raoof and Kaushik 2002).

Strains of *Limnospira* are some of the fastest known photosynthetic organisms, with doubling biomass times shorter than a day under optimum conditions (Shi et al. 2016). Which benefits from high RuBisCO content and efficient carbon-concentrating mechanisms (Singh and Gaur 2026), that benefits from occurring naturally at bicarbonate-rich environments. In addition, high performance has been attributed to the extremely high tolerance to oxygen supersaturation (>300% dissolved O_2_), conditions where most cyanobacteria suffer from photorespiratory stress (Marquez et al. 1995; Franke et al. 2022).

Industrial strains, like PCC 7345 (Paramanya et al. 2024), Paracas R14(Reichert et al. 2006) and SAG 85.79 (Ljubic et al. 2018) have been widely studied in growth-rate, nutrient-optimization, and bioreactor performance. These strains have decades of cultivation history and predictable behavior. Because domesticated strains have undergone unintentional selection (Benedetti et al. 2018), genomic streamlining, adaptation to high-alkalinity, high-light, nutrient-rich conditions, our current understanding of *Limnospira* likely underestimates the ecological plasticity of wild populations (Cohen 1997; Sili et al. 2012).

Domesticated strains only represent a narrow, human-selected subset of industrially relevant traits, while wild strains capture the ecological, physiological, and evolutionary diversity of *L. platensis.* Therefore, the study of wild strains is necessary to understand natural adaptation in the context of its evolutionary history.

In this work, we present comparative growth of the five spirulina strains at different temperature and light growth conditions to identify differences under a single variable in controlled conditions. Then, we compared their biochemical and biophysical performance under a naturally fluctuating light environment to explore the architecture diversity of energy utilization in these strains and elucidate if this diversity correlates with phylogenetic relationships.

We selected UTEX strains: LB 1926, LB 1928, LB 2340, LB 2342 and 3086. Strains LB 1926 and 1928, are American oceanic strains, while LB 2340, LB 2342 and 3086 are from African salt lakes (UTEX 2026; Starr 1978). Strain LB1926 (equivalent to PCC 7345 and deposited by F.T. Haxo in 1972 at UTEX) is the most studied of the selected strains, although, the original strain was LB 1928 (isolated by Lewin, 1969). LB 2342 is the only strain included in this work that has been originally classified as *L. maxima*, and sub-strains (JN831262 and Lefevre 1963/m-132 FJ798616) seemed to have diverged significantly in a short period of time (Nowicka-Krawczyk et al. 2019). From the strains selected in this work, UTEX 3086 is one of the two fully sequenced strains of *L. platensis* (Shiraishi and Nishida 2023).

## 2. Materials and Methods

### 2.1. Biological Material

Five Spirulina strains were obtained from the UTEX Culture Collection (Houston, Texas United States of America). A volume of 1 mL of strains LB 1926, LB 1928, LB 2340, 3086 and LB 2342 were transferred individually from its original tube into a disposable Roux bottle (T-25, 40 ml max. vol.) containing 10 mL of the recommended UTEX medium to prevent further shock. Strains 3086, LB 2340, and LB 2342 were maintained in UTEX Spirulina medium, whereas strains LB 1926 and LB 1928 were maintained in UTEX Enriched Seawater. Media was purchased directly from the UTEX collection to avoid any nutrient shock. Table 1 summarizes the information available for the strains used in the present work. Samples were kept at 23°C with a photoperiod of 8 hours of light with an intensity of 50 µmol m^-2^ s^-1^ using neutral white fluorescent light tubes (standard storage conditions at the Brock University’s Microphyte Collection, Ontario, Canada).

**Table 1:** Information about the origin of strains and equivalence with other commonly studied strains.

| UTEX ID | Origin | Species | Isolation | Equivalence | Original collection |
| --- | --- | --- | --- | --- | --- |
| LB 1926 | Del Mar Slough, San Diego, CA, USA | platensis | R.A. Lewin (1969) | UTEX LB 1928<br>PCC 7345<br>ATCC 29408 | F.T. Haxo (1972) |
| LB 1928 | Del Mar Slough, San Diego, CA, USA | platensis | R.A. Lewin | UTEX LB 1926<br>PCC 7345 | UTEX |
| LB 2342 | Natron Lake, Tanzania | maxima | M. Lefevre (1963) | CCAP 1475/9<br>SAG B 84.79<br>CS-328 | SAG |
| 3086 | Lake Chad, Chad | platensis |  | NIES-39<br>IAM M-135 | NIES |
| LB 2340 | Natron Lake, Tanzania | platensis | G. Laporte (1963) | SAG B 85.79<br>NIVA CYA 120 | SAG |

After two weeks, copies of each strain were transferred to Zarrouk media (Phytotreat, the Netherlands) to standardize growth conditions. Pre-made powdered media was dissolved in distilled water and autoclaved before use. Before any experimental procedure, samples were adapted to Zarrouk media for at least two weeks and re-cultured twice to ensure a stable phenotype was obtained.

### 2.2 Growth measurements

Determination of growth was done using optical density (OD) via optical spectroscopy using Zarrouk media as a blank. OD measurements were calibrated against biomass density (see Figure S1 and Table S1). OD measurements were taken at 750 and 850 nm using a modified near- infrared–visible spectrometer (SPECTRONIC™ 200 Visible Spectrophotometer. Thermo Fisher Scientific, Waltham, MA, USA) capable of measuring 10 mm (optical path) cuvettes, cylindrical tubes and T25 Roux bottles. The modified sample holder designed and 3D-printed by Dr. Zavafer can be downloaded from our website (biophotonicsbrock · GitHub) and is available as a 3D printing file (STL) or as source file (Trimble Inc. SketchUp, Version 2024 [Computer software]. https://www.sketchup.com).

Biomass determinations were done by filtrating 100 mL of culture using a Whatman No. 8 paper filter, which was washed twice with 200 mL of distilled water. Filtration was followed by drying the paper in an oven at 60 °C for two days. To determine biomass density, measurements of the weight before and after filtration were used to estimate the total amount of biomass and divided against the 100 mL.

### 2.3 Growth analytics

To determine the maximum growth rate (*r_max_*), we applied an algorithm based on (Sekse et al. 2012) using spline calculated curves. OD measurements were plotted against time, and a spline model was applied to each time series (each series is composed by the time course of individual samples) to obtain a continuous data series. Then the first derivative of each resultant spline was calculated for each time series. Because the maximum value of the first derivative represents the maximum growth rate, this value was recorded for all curves and averaged between samples in the same treatment. Then maximum growth rates were converted in doubling time by using the ratio *ln*2/*r_max_*.

### 2.4 Growth conditions in controlled environments

Two environmental stressors were examined in this study: temperature and photon flux density (PFD). For both experimental series, each strain was pre-adapted to the target condition for one week to ensure that a stable physiological phenotype was established. The duration of each growth cycle was defined by the point at which cultures surpassed their maximum growth rate (see Growth analytics section).

For temperature tolerance, cultures were transferred to cylindrical glass test tubes (15mL, Fisher Scientific). Five temperatures (13, 23, 26, 30, and 33°C) were tested to determine the optimal temperature for growth of each strain by growing the sample at either incubators (NAPCO and Conviron) or heating blocks (Fisher scientific). Each culture had a volume of 6.5 mL and came from mixing alga from an acclimated culture in the temperature. The ratio of the mixture depended on the OD of the culture, as the aim was to standardize all samples to a similar low initial OD.

Samples at each temperature were placed in an aluminum tray filled with glass beads. This allowed better thermal contact while keeping uniform light diffusion. The tray was positioned 15.8 cm below a generic neutral white (5000 K) LED arrays. Position of each tube was randomized daily to avoid the effect of light hotspots. Each LED was powered by an adjustable benchtop laboratory power supply (Nawei sz, China) and current was adjusted to produce an average PFD of 55 µmol m^-2^ s^-1^ photons at the sample level. Light cycle had a duration of 8 hours per day.

To test light tolerance, a culture with low concentration (OD_850_ < 0.10) of 240 mL in Zarrouk medium was prepared for each strain and subsequently divided into six 40 mL cultures in a small Roux (T-25) bottles, each containing an autoclaved magnetic stirring bar. Afterwards, each bottle was stirred individually (Thomas Model 14) and illuminated with a generic 50 W COB LED light with a square active area of 35 mm^2^. Light intensity was measured using a quantum meter (Spot-On, United States of America). Current limiting was established using high power resistors (> 1 W) to obtain different PFD values and an adjustable benchtop laboratory power supply (Nawei SZ) was used to drive the LEDs. An approximate 30 rpm was maintained in each bottle to ensure sample homogeneity. The applied PFD of each bottle ranged from 12 to 350 µmol m^-2^ s^-1^ using an 8 h photoperiod. The optical density was measured directly in the culture bottle to monitor overall culture status, particularly during the initial acclimation phase. Also, an aliquot of each culture was transferred to a cuvette with a 0.5 cm path length to obtain more accurate measurements as cell density increased and to enable consistent long-term tracking of growth.

It is worth noting that LB 2340 was not included in this experiment as their cells have a tendency of individual cells to aggregate into clusters rather than remaining evenly dispersed in suspension, and required mechanical stirring to form a homogenous solution, which is necessary to measure OD.

### 2.5 Greenhouse growth

Once optimum temperature was established, cultures of each strain were transferred to a greenhouse for culturing. This was carried out between the months of November 2025 to May 2026 at the St. Catharines campus of Brock University, ON, Canada (43.1190° N, 79.2490° W). Samples pre-adapted to 30°C were scaled up to 400 mL cultures Roux bottles (T125) and submerged in a generic glass fish tank equipped with a ceramic heater and a thermostat. The temperature of the water was set to be 30°C. Initial attempts to establish the culture in the absence of mechanical stirring or air bubbling were unsuccessful. Thus, Roux bottles were equipped with bubblers made from autoclaved silicone tubing interfaced to the bottle via carbon-fibre, autoclave- compatible, 3D printed caps. These caps were also equipped with an exhaust port for gas exchange using a sterile filter tip (250 µL) to avoid contaminants going into the culture. Bubbling was done using a small air compressor coupled to air humidifiers using distilled water (to avoid excessive evaporation). This setup resulted in stable growth for 6 months. Harvesting of culture and addition of fresh media was done weekly by diluting each Roux bottle by half its volume.

### 2.6 Protein Quantification

Greenhouse adapted cultures of all strains were used for protein and phycocyanin determination. Estimation of the biomass was done using OD measurements on one aliquot of 1 mL on the day of measurements before protein analysis was carried out, and biomass was calculated based on pre-established calibration curves.

Protein determination was done using a Bradford assay. A sample of 50 mL was extracted from each bottle in the greenhouse setup. Then, 35 mL of the sample was placed into a falcon tube of 50 mL. Continuing, the samples were sonicated (U.S. Solid, U Cleveland, Ohio) at 20kHz with a 20% amplitude for 7 minutes with a pulse on for 25 seconds and pulse off for 20 seconds. Afterwards, a Bradford assay was performed using the Quick Start™ Bradford Protein Assay (Bio-Rad) with bovine γ-globulin as the standard. To start, the standard curve preparation began by adding 100 µL of each protein standard (from 0.125 mg/mL to 2.00 mg/mL) to 5 mL of Coomassie Brilliant Blue G-250 dye reagent.

Following dye addition, spectroscopic measurements were performed at 595 nm for both the standards and the samples. Then the standard calibration curve was generated by plotting OD at 595nm against protein concentration of the standards and protein concentration was estimated as per manufacturer indications. The calculated protein concentrations were subsequently used to determine the total protein content and its percentage relative to the total dry weight for each strain. Spectroscopic determinations were done using a Thermo Scientific Genesys 150 spectrophotometer (Thermo Fisher Scientific, Waltham, MA, USA).

### 2.7. Phycocyanin extraction and determination

A sample of 100 mL was extracted from each bottle in the greenhouse setup. Following, 1 mL was separated from each sample to measure the OD of the respective culture. Then, another part of the sample was used to fill Falcon tubes (volume 50 mL) for each strain with a volume of 30 mL. Then, the samples in the Falcon tubes were centrifuged at 4500 G for 30 minutes using a Eppendorf 5804R centrifuge to separate the alga from the media, after discarding the media (supernatant), and the pellet was resuspended using 25 mL of an ammonium sulfate solution with a concentration of 1.1 M. Afterwards, the samples were sonicated at 20kHz with a 20% amplitude for 7 minutes with a pulse on for 25 seconds and pulse off for 20 seconds. After sonication the samples were centrifuged for 15 minutes at 5,000 rpm, then the supernatant was discarded, and the pellet was resuspended in a solution of CaCl2 with a concentration of 0.1M. The samples were centrifuged again for 15 minutes at 5,000 rpm. Spectroscopic analysis was performed in the supernatant of the samples to calculate the concentration of C-Phycocyanin (PC) and A-Allophycocyanin (APC) in the samples using OD at 615 and 652nm (measured in a Thermo Scientific Genesys 150 spectrophotometer) and the following equations:

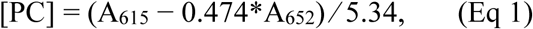

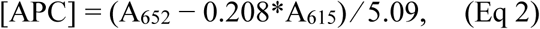

Finally, the concentration values were used to calculate the percentage of weight that Phycocyanin represents in each specific strain and phycocyanin determination. Estimation of the biomass was done using OD measurements on one aliquot of 1 mL on a measurement day before protein analysis was carried out, and biomass was calculated based on pre-established calibration curves.

### 2.8 Radiometry

Radiometric determinations (Photon Flux Density and daily light integral) were done using a SpotOn Quantum PAR Light Meter (Innoquest Inc., Woodstock, IL, USA) as outlined in Zavafer et al. (2023). To match light conditions at the UTEX collection, a portable spectroradiometer (Photon Systems Instruments, PSI. SpectraPen spectroradiometer. Drásov, Czech Republic.) was used where equivalent lux were transformed into PFD units and the light intensity adjusted to it.

To determine the daily light integral (DLI) inside the greenhouse, daily values of the DLI were obtained from the monthly published report by the Vineland Research and Innovation Centre (VRIC) in Vineland, Ontario (nearest station for radiometry) and are accessible at the website (ongreenhousevegetables.ca). Daily values were adjusted to the attenuation loss by the greenhouse panels measured by determining the average PFD w inside and outside of the greenhouse at noon. An average attenuation of 30% was used to adjust daily DLI inside the greenhouse.

Greenhouse experiments had an average DLI of 10.5, 7.05, 9.77, 15.18 and 16.72 mol m^-2^ s^-1^ for the months of November 2025 to March 2026. The maximum DLI was 22.75 mol m^-2^ s^-1^ and the lowest 1.4 mol m^-2^ s^-1^. From the 133 recorded days of the experiment, 73% of the days of the experiment L*imnospira* cultures were exposed to DLI values > 10 mol m^-2^ s^-1^ (maximum DLI used in our experiments under controlled conditions). Based on these radiometric conditions, the greenhouse was considered a high light environment.

### 2.9 Biophysical characterization via chlorophyll-a fluorometry

Greenhouse adapted cultures of all strains were used for chlorophyll *a* fluorometry. Dark adaptation was measured using the Kautsky induction curve (also known as OJIP curve) using an OpenJIP fluorometer equipped with a 450 nm COB 10 W LED and illuminated with 5,000 μmol m^-2^ s^-1^ for 3 s (Bates et al. 2019). Samples were dark adapted with 1, 5, 15 and 30 minutes and the ratio F_V_/F_M_ was calculated.

A rapid light curve protocol was applied to samples using a Water PAM (Walz GmbH. (n.d.), Effeltrich, Germany) using the Ralph and Gademann (2005) approach to determine the relative electron transport rates.

To parametrize the decay after the *F_M_* during Kautsky induction curves we used an empirical sigmoidal model:

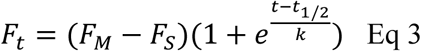

where *F_t_* is the fluorescence at a given time, *F_S_* is the fluorescence at the S step, t is time, *t*_1/2_ is the half-time and *k* is a time-scaling parameter shape parameter (in time units). *k* controls how wide the logistic transition is. Since multiple strains reach *F_M_* at different times, we normalized the time domain to the fluorescence value where *F_M_* was reached (i.e., *t_Fm_* = 0).

### 2.10 Phylogenetic analysis

To compare the phylogenetic relation between strains we subset phylogenies of three independent works: Nowicka-Krawczyk et al. (2019); Roussel et al. (2023); Pinchart et al. (2024) to include only the species of interest. Here we assume that UTEX strains were equivalent to the strains used as per Table 1 and as per information at the UTEX database.

### 2.11 Statistical data analysis

For the statistical analysis of the results of these experiments the analytical software Excel, Anaconda Python, and Origin Lab 2023b were employed.

## 3. Results

We compared the phylogenetic relationships among the strains of interest across four phylogenetic trees constructed from different genomic regions and systematic methods. Three of the four trees supported the grouping of UTEX LB 2340 and LB 2342 into a clade, while UTEX 3086 and LB 1928 consistently formed another clade. The only contrasting tree was the one presented in Figure 1a where multiple sub strains of LB 2342 act as an outgroup to the clade formed by UTEX LB 1928, LB 2340, and another LB 2342 strain. Also, this tree had multiple polytomies and was not considered further in this study. UTEX LB 1926 was not included in any phylogenetic tree.

**Figure 1.**
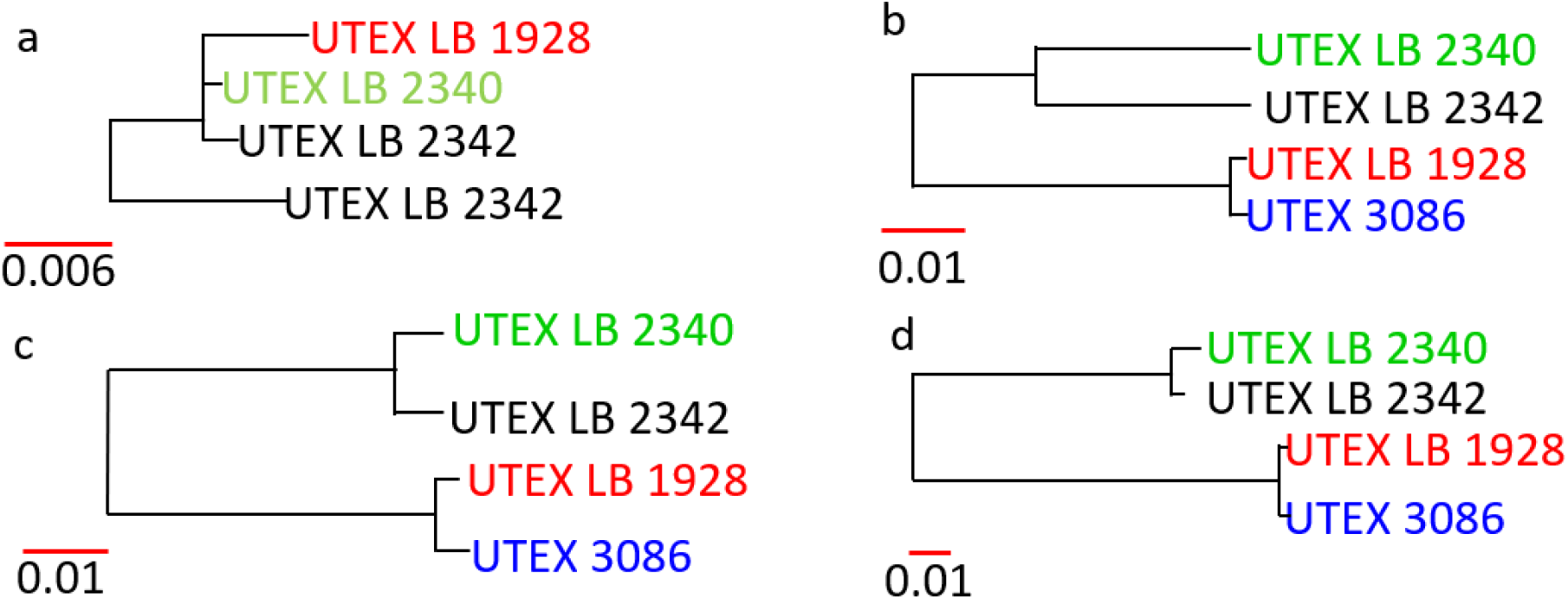
Comparison of different phylogenies of *Limnospira platensis*. (a) Phylogeny according to Nowicka-Krawczyk et al. (2019) based on 1800 bp fragment of the gene for 16S rRNA. (b) Phylogeny according to Roussel et al. (2023) based on 16S-ITS concatenated sequences. (c) Also, Roussel et al. (2023) but consensus phylogenetic tree based on the 2672 most conserved genes. (d) Phylogenomic maximum likelihood tree from Pinchart et al. (2024) of three newly established Limnospira Metagenome-Assembled Genomes (MAGs) from spirulina cultures.

To evaluate whether the phylogenetic relationships observed among the strains correspond to similarities in their growth under different environmental conditions, we examined two major growth stressors: temperature and light intensity (Vonshak 1997). Figure 2 summarizes growth performance across the tested temperature range. Although all four strains shared the same optimum temperature of 30 °C, their maximum growth rates (*r_max_*) differed significantly. All strains also exhibited an asymmetric temperature tolerance, showing a broader range of growth in the colder temperatures than toward warmer ones. This is a common feature in mesophilic cyanobacteria (Huesemann et al. 2023). While growth was detectable at 13 °C, the actual doubling time exceeded 10 days for all strains (Figure 2b includes average doubling times for context).

**Figure 2:**
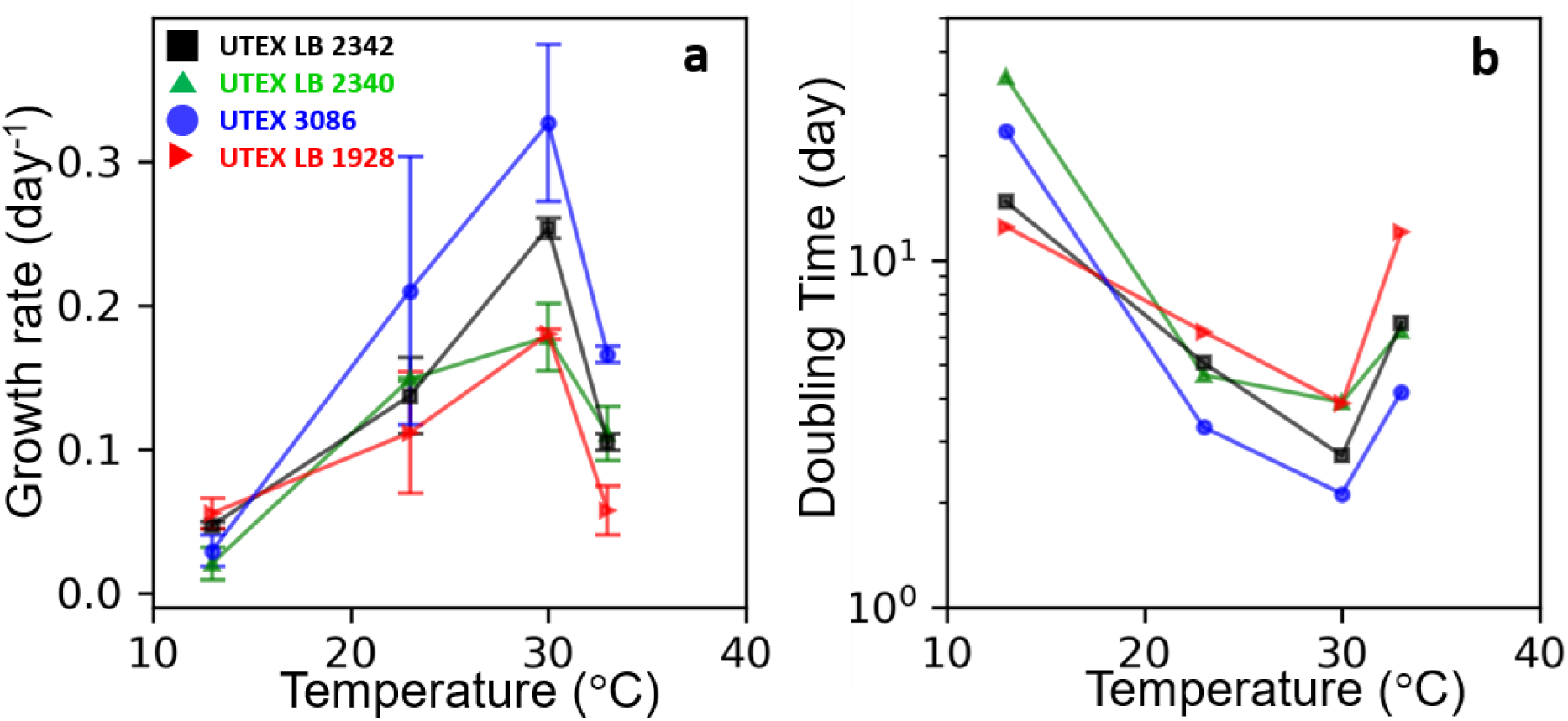
Effect of temperature in the maximum growth rate of different strains. (a) Data presented as a growth rate (d^-1^) each point represents the mean ± std of n = 4 samples. (b) Equivalent doubling time calculated for each average growth rate. No std deviation was included for ease in the visualization as doubling time is a logarithmic parameter.

UTEX 3086 presented the highest rate (∼ 0.3 day^-1^) followed by UTEX 2342 (∼ 0.25 day^-1^) while both UTEX LB 2340 and 1928 presented rates of ∼ 0.15 day^-1^ (Figure 2a), which translates into doubling times of 1, 1.5 and 3 days respectively (Figure 2b). No correlation between phylogenetic relationship and temperature was found in this test.

There is a possibility that the observed growth in our temperature experiment was hindered by the lack of stirring and the use of a low photon flux density (PFD, 50 µmol m⁻² s⁻¹; equivalent to a daily light integral, DLI, of 1.44 mol m⁻² d⁻¹). To address these limitations, we tested growth under vigorous stirring across a range of PFD values (Figure 3). In all strains, growth followed a light-response curve consistent with the Platt model (Figure 3a), with each strain exhibiting a distinctive maximum photosynthetic capacity (Pₘ, Figure 3b) and efficient light use at low irradiance (α, Figure 3c). Note that, instead of presenting data in terms of PFD, Figure 3a displays values as DLI to facilitate comparison with greenhouse conditions presented further on in this study.

**Figure 3:**
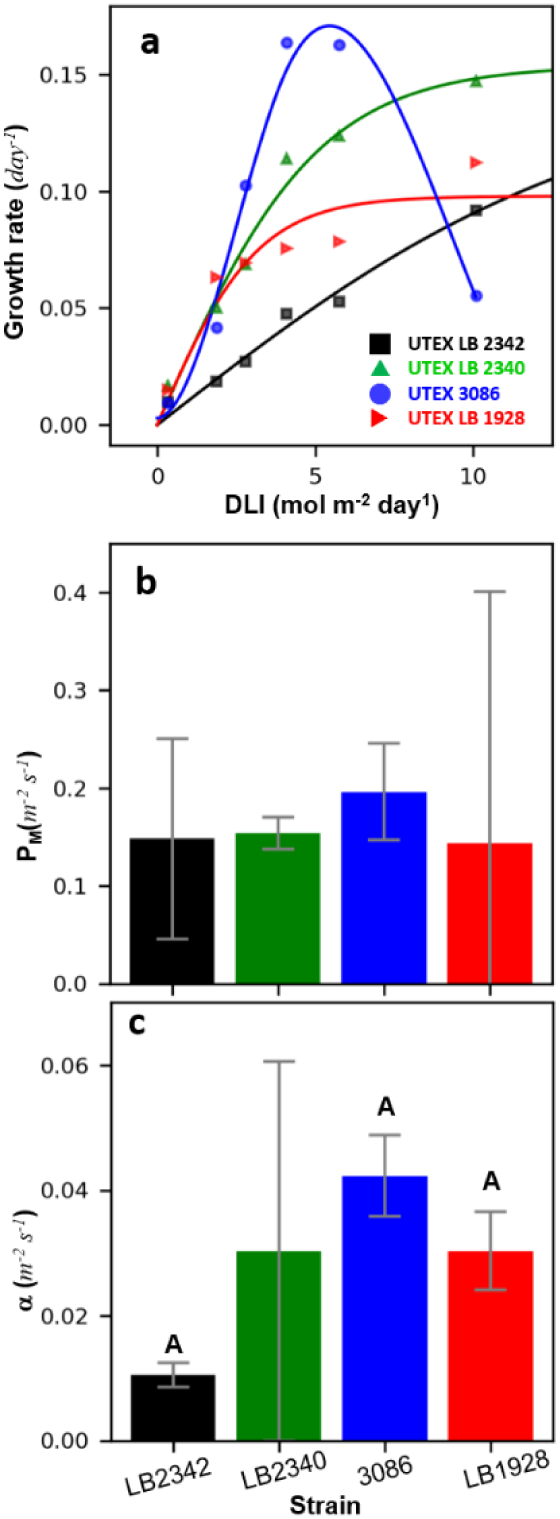
Comparison of light tolerance between different strains. (a) Growth versus light intensity (displayed as DLI values), a characteristic experiment is displayed as example. (b,c) Fit parameters (PM and α) to the Platt model (or spline Platt model) ± standard error of the fit, p < 0.05 is used to determine significance. Character “A” at the top of bars indicate that values where significantly different to those observed for LB 2342 only, absence of such characters indicate no significant difference.

Similar to our temperature observations, UTEX 3086 showed the highest apparent growth rate. However, when Pₘ was estimated, no significant differences in maximum photosynthetic capacity were detected among strains (Figure 3b). What did differ was the DLI at which each strain reached Pₘ (Figure 3a). Strikingly, UTEX 3086 was the only strain that exhibited photosensitivity, with clear photoinhibition observed at DLI values exceeding 5 mol m⁻² d⁻¹ (corresponding to ∼200 µmol m⁻² s⁻¹, see Figure 3a). It is also worth to note that the observed *r_max_* was half of that observed for the temperature experiment. This can be explained from the fact that this experiment was done at 23°C and difference in the actual optical path. Since this experiment was done using a longer optical path to those used in the temperature experiment, since shorter paths decrease the effect of self shading (tubes of 10 mm diameter, versus 28 mm optical path in the Roux bottles), a bio-optical effect could also be responsible for the observed difference between experiments (Serôdio and Campbell 2021).

Regarding α, the initial slope, correlating with the quantum yield of PSII electron transport (Saroussi and Beer 2007), LB 2342 was significantly lower, and thus less efficient, than UTEX 3086 and LB 1928, while LB 2340 showed no significant difference due to poor curve fitting (Figure 3c). Under our experimental conditions, LB 2340 could not be saturated, resulting in a poor fit. UTEX 3086 appeared to have a higher α compared with LB 1928, but these differences were not statistically significant.

From the comparison of the two environmental variables, temperature and light, the two traits with the most significant differences were *r_max_* and light tolerance. Thus, we compared the photophysiology of the strains under a fluctuating light environment inside a greenhouse to determine if the observed differences were relevant in a more natural environment. Since all strains had the same optimum temperature, cultures were grown at 30°C to reduce complexity of the study. First, we compared the fraction of total biomass that is only protein between strains (Figure 4a), where it was observed that the range of protein is found to be between 57 to 70%, which is consistent with literature values (Ogbonda et al. 2007). No significant difference was found between strains. Then we analyzed the content of C-phycocyanin, where the range was between 7.5 to 15% of the total biomass among different strains (Figure 4b). Only strain LB 1926 and 1928 showed significant difference, where the former was 60% higher. Finally, for comparison, we determined the fraction of total protein which is C-phycocyanin finding a range between 15 to 25% among different strains, but only strains LB 1926 and 1928 showed significant difference (Figure 4c).

**Figure 4:**
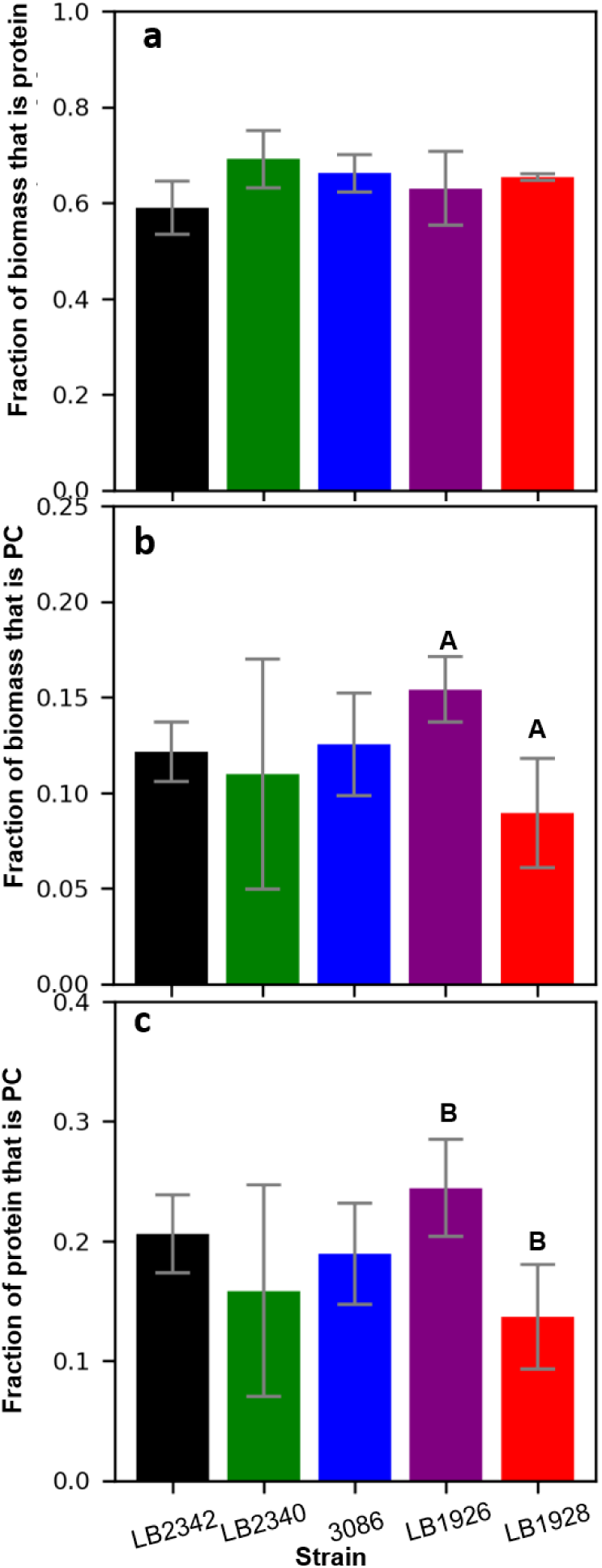
Protein and C-phycocyanin productions in spirulina strains. (a) Fraction of total biomass that is protein measured via a Bradford assay. (b) Fraction of total biomass that is C-phycocyanin of the total biomass. (c) Fraction of total protein that is C-phycocyanin. Values correspond to the mean of two different experiments each with 4 technical replicates (4 aliquots of the same sample) measured on each day of the experiment for strain ± standard error.

Protein analysis above indicates that despite differences in growth rate and light tolerance, the difference in these two traits cannot be attributed to bulk protein expression. Thus, a possible explanation to the observed differences in growth could be caused by fine tuning of photosynthetic reactions(Rochaix 2013; Pribil et al. 2018; Gan et al. 2019), and not general metabolic differences.

We examined the maximum value of F_V_/F_M_ (an indicator of the functional state of PSII charge separation, Sipka et al. (2021)) under different durations of dark adaptation. This test would inform how efficiently PSII reaction centers recover after being exposed to light (Figure 5a). As it is the case in cyanobacteria, maximum values of F_V_/F_M_ values ranged from 0.32 to 0.5, and most strains displayed their maximum value after only 1 min of dark adaptation, with the exception of UTEX LB 2340 which perform slightly better at 5 min. Also, like is the case of cyanobacteria and microalgae, prolonged dark adaptation yielded lower F_V_/F_M_. We also found that UTEX LB 2340 was the best performer with a maximum F_V_/F_M_ of ∼0.52, while UTEX LB 2342 and LB 1926 performed almost identically. Finally, UTEX LB 1928 and 3086 displayed the lowest values of F_V_/F_M_, being this latter strain the worst performer. UTEX LB 2340 was the most tolerant and UTEX 3086 the most sensitive, and these results correlates well with the light tolerance observed in Figure 3, as DLI in the greenhouse ranged between 5 to 30 mol m⁻² d⁻¹, which can be considered high light for all strains.

**Figure 5:**
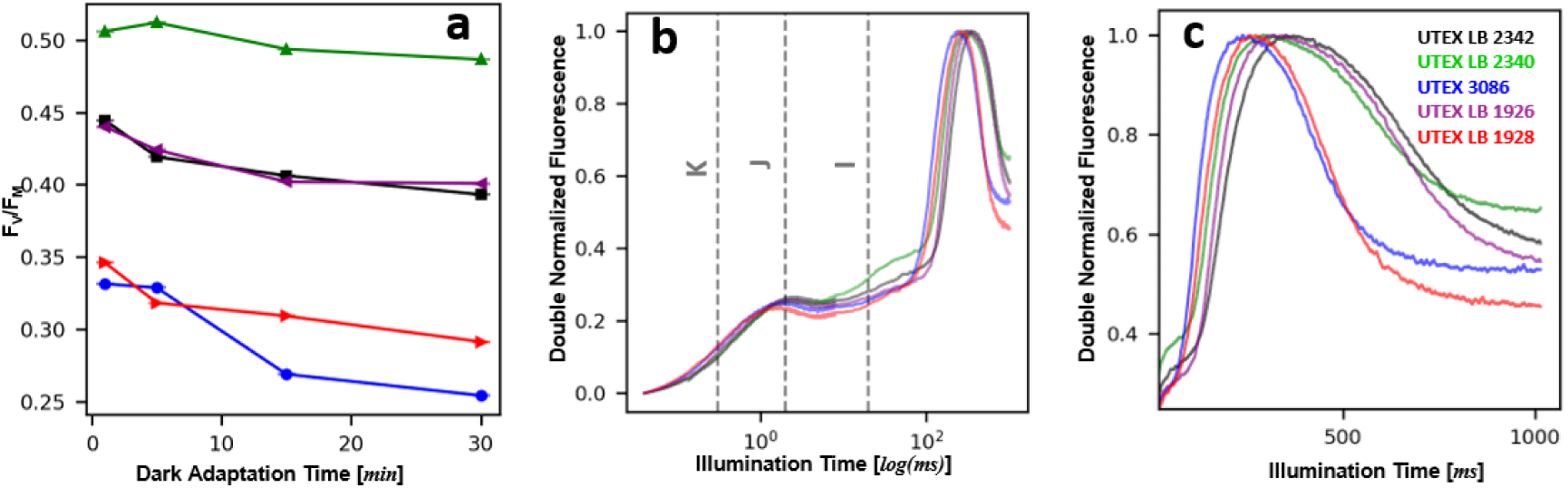
Biophysical determinations of the chlorophyll *a* fluorescence measured via the Kautsky induction curve. (a) FV/FM values as a function of dark adaptation time. (b) Kautsky curves double normalized (0 = FO and 1 = FM) to compare morphology of the curve, data presented in logarithmic scale. (c) Decay after FM in the Kautsky induction curve presented in linear scale from the I-step. Panels b,c present only average curves to ease visualization.

Since light tolerance can be explained in changes in the organization of the components of the thylakoid membrane, we examined the shape of the of Kautsky induction curve (OJIP curves) which can be used as an *in-situ* indicator of such changes (Lage-Pinto et al. 2008). We observed (Figure 5b) that although all strains share similar shapes, all displaying a sigmoidal OJ phase (phase linked to reduction of Q_A_), a quenched JI phase (phase linked to electron transport between Q_A_ and Q_B_/PQ) and a steeped rise of the IP phase (linked to PSI and other accessory processes, Toth et al. (2007)). In fact, the most striking difference between curves was the time to reach F_M_ and the fluorescence decay after it (Figure 5c), where UTEX LB 2342, LB 2340 and LB 1926 had similarities, such as slower decay of fluorescence and time to reach F_M_. Meanwhile, UTEX 3086 and UTEX LB 1928 displayed faster quenching kinetics.

Because F_M_ corresponds to near to full reduction of all active PSII reaction centers, the decay of F_M_ is linked to two main factors (Zhu et al. 2013; Vredenberg 2018; Grigoryeva 2020): (1) electron transport in light adapted state; (2) the onset of regulatory/quencher processes (e.g., non-photochemical quenching or state transitions). To determine if these changes were caused by electron transport, we fit this decay to a sigmoidal model and compare it against rates of electron transport under a light adapted state (Figure 6).

**Figure 6:**
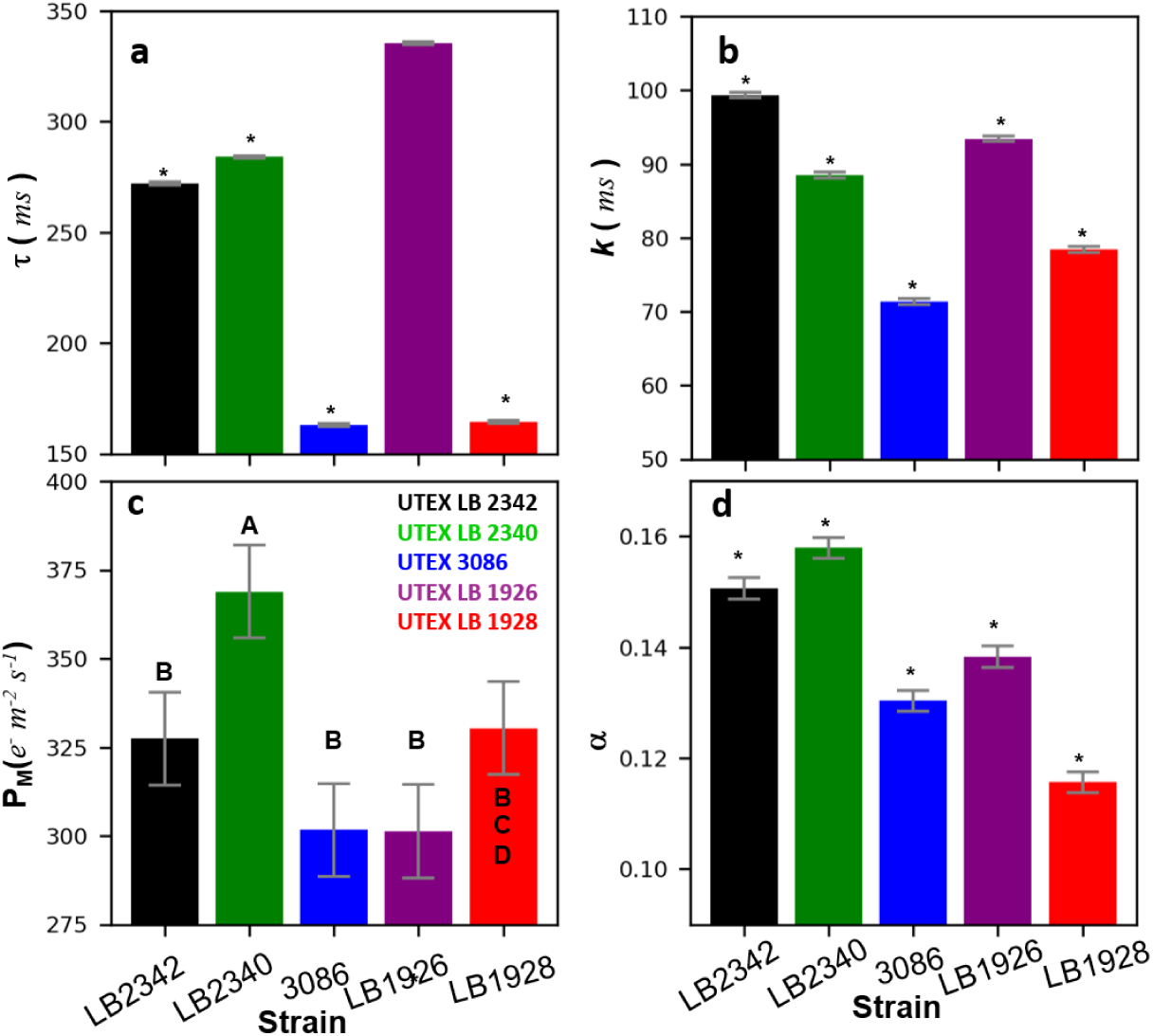
Comparison between decay of the fluorescence level after FM in the Kautsky induction curve versus the maximum photosynthetic rate (PM) and α obtained from rapid light curves. (a) *t*_0_ (onset/inflection time), (b) Half-decay time (*t*_1/2_) of the Kautsky induction curve, (c) PM and (d)α. Data correspond to the fit value ± standard error calculated from experimental curves (3 < n < 4). Statistical significance was estimated p > 0.5 and characters in the top indicate significant difference among all strains (*) panels (a,b,d) and for panel c: significant different to LB 2342 (A), LB 2340 (B), LB 3086 (C) and LB 1928.

The sigmoidal model provides two parameters *t*_0_ (onset/inflection time) and Half-decay time **(*t*_1_**_/**2**_) and there are displayed for every strain in Figure 6ab. *t*_0_ indicate how quickly the decay switches from slow to fast and marks when the dominant quenching process drives the decay, while *t*_1/2_ is a proxy for the rate at which PSII reaction centers reopen and/or quenching builds. All strains had significantly different values of both parameters, being *t*_0_ (Figure 6a) the slowest in UTEX LB1926 and the quickest UTEX 3086 and LB1928. This pattern was similar to *t*_1/2_ Figure 6b, the only behavior difference in this parameter was between LB 2340 and 2342 where *t*_1/2_ was slower for the latter.

The Pm and α for relative electron transport rates (Figure 6cd) showed that only UTEX LB 2340 had a higher P_M_ than other strains Figure 6a. Meanwhile, α was significantly different among all strains, where UTEX LB 2340 was the highest, followed by strains LB 2342, LB 1926, 3086 and LB 1928. Finally, we found a close similarity in the behavior of *t*_1/2_ and α where all strains behaved very similarly, suggesting that the empirical parameter *t*_1/2_ correlates well with the α (linked to the quantum yield of PSII electron transport), and indicating that the halftime decay is mostly explain by efficiency of electron transport in the light acclimated state of photosynthesis.

To conclude our experimental measurements, we compared the morphology among strains under the microscope to see if any changes in the length or shape of the trichomes can be seen between strains and environments, as stress determines morphology of trichomes (Ma and Gao 2009). A comparison of cultures in controlled environment (DLI: 1.44 mol m⁻² d⁻¹, 30 °C) versus the greenhouse is presented in Figure 7. UTEX LB 2340 (Figure 7ab) displayed a straight trichome colonies, while the rest of strains displayed spiral trichomes (Figure 7c-j). UTEX LB 2342 had shorter trichomes (Figure 7cd), than the other spiral strains. Finally, spiral strains presented slightly longer trichomes under controlled environments (Figure 7a,c,e,g,i) than in the greenhouse (Figure 7b,d,f,h,j). UTEX 3086, LB 1926 and LB 1928 expressed very similar phenotypes.

**Figure 7:**
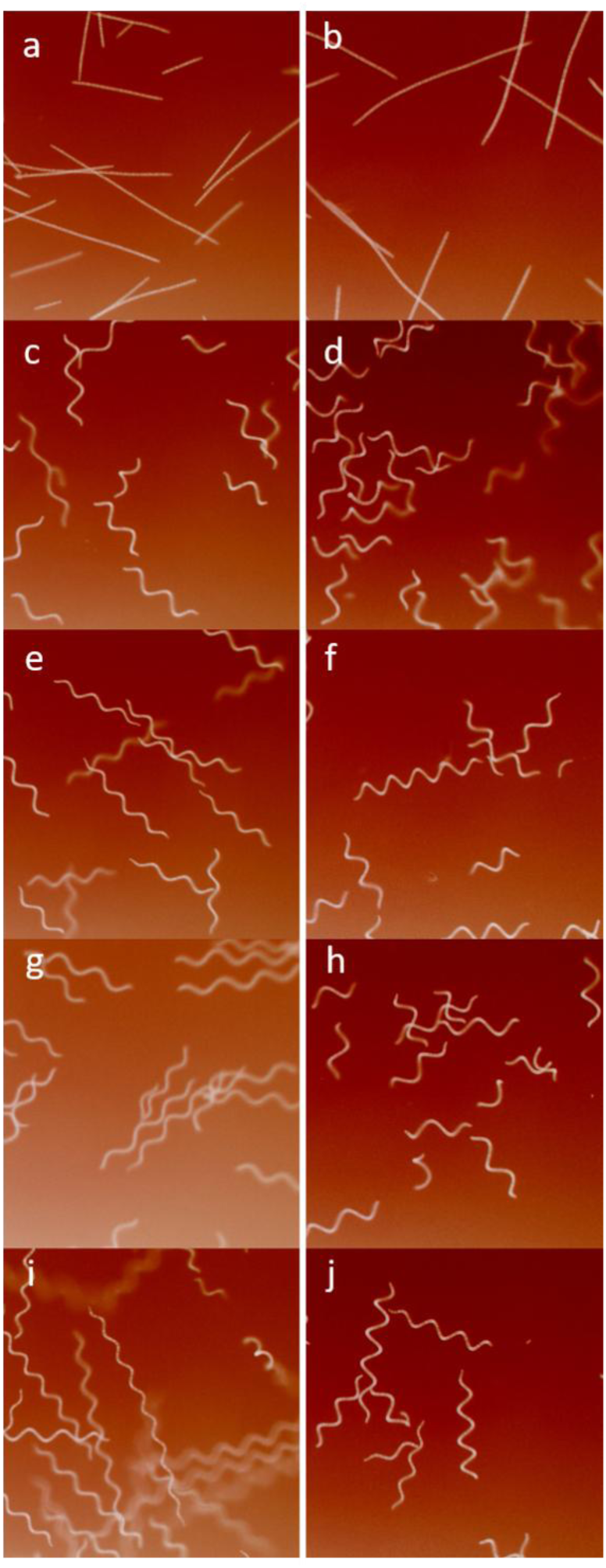
Morphological comparison between strains grown in a controlled environment (a, c, e, and, i) versus the greenhouse (b, d, f, h and j). (a,b) UTEX LB 2340, (c,d) LB 2342, (e,f) 3086, (g,h) LB 1926 and (h,i) LB 1928. A characteristic image is presented for each strain and condition. To enhance contrast a false color image is used over a composite image composed by the green and red channels.

## 4. Discussion

In this study, we evaluated growth-related phenotypic traits in the context of phylogenetic relationships among multiple *L. platensis* strains under different environments. Some traits were consistent within clades, while others varied markedly despite how closely phylogenetically strains were. For example, strain UTEX LB 1926 and 1928 (according to UTEX records) were at some point the same isolate, but here we show that both performed very differently, and LB 1926 performed more closely to UTEX LB 2342. Other traits, however, such as net protein content and C-phycocyanin appeared to be very conserved across the different strains within *L. platensis* and we did not observe any statistical difference during the study.

Based on three phylogenies from two independent studies (Figure 1), the strains studied here arranged into two clades: clade A (UTEX LB 2340 and LB 2342), and clade B (UTEX 3086 and LB 1928). The formation of a clade A is not surprising since both strains were isolated from the same geographical region (see Table 1). However, it was surprising to find that UTEX 3086 and LB 1928 have a closer phylogenetic relationship (Roussel et al. 2023; Pinchart et al. 2024) with an American strain from the Pacific coast rather than other African strains.

Upon examination, some traits such as F_V_/F_M_ (Figure 5a), decay after F_M_ (Figure 5c and 6ab) and electron transport in light adapted state (α and P_M_, Figure 6cd), had similar values among members of the same clade. Thus, traits associated to fine tuning in photosynthetic reactions performed similarly in phylogenetic related strains. In contrast, traits associated with growth such as *r_max_* at different temperatures and light tolerances showed no clear phylogenetic relationship. This bears the question how relevant biophysical traits measured here are in the actual context of algal growth for spirulina.

For example, photosensitivity of UTEX 3086 is explained from biophysical data, as this strain presented hindered growth at high light intensity (Figure 3), and also exhibited the lowest F_V_/F_M_ in the greenhouse (a high light environment). Another example is LB 2342 which withstand better high light and is not photoinhibited under the experimental conditions (Figure 3) and had a higher F_V_/F_M_ value when compared against UTEX 3086 (Figure 5). Typically, lower values of F_V_/F_M_ in the context of light intensity are associated with sustained quenching mechanisms as a response to excessive excitation by PSII (Vass and Cser 2009; Serôdio and Campbell 2021; Zavafer 2021). While not all biophysical measurements can explain growth, they allow us to understand the behavior of some of the observed strains.

However, attributing high or low values of *r_max_* cannot be used directly to explain differences in electron transport, as no direct correlation was found between growth as a function of light intensity (Figure 3) versus values observed for the rapid light curves (*i.e.,* rates of electron transport, Figure 6). For example, rapid light curves measure non-steady-state artifacts, and there may be underestimations of maximum electron transport rates due to the short duration of each light step (Huang et al. 2021). Thus, other factors not measured in this work such as the carbon fixing and concentrating mechanisms (Cheng et al. 2017; Gan et al. 2019; Singh and Gaur 2026), respiration (Berry et al. 2003), among others may play a more significant role on the growth rates.

Total protein content and fluorescence measurements obtained from greenhouse samples do not directly translate to the results from controlled-condition experiments. A fluctuating light environment imposes physiological challenges on cyanobacteria that differ substantially from those experienced under constant PFD conditions (Andersson et al. 2019). Radiometric measurements confirmed that the greenhouse DLI was considerably higher than the values used in the light-tolerance experiment. Even so, the light-tolerance assays provided a useful reference point, allowing us to evaluate whether the greenhouse environment exceeded or fell below the optimum irradiance range for each strain.

A factor that can explain the differences in growth is how quickly Limnospira strains have evolved in controlled environments. For example, UTEX LB1926 and LB1928 have been maintained separately since 1968, and the isolation for 58 years caused the two strains to accumulate enough mutations to develop different traits. Traits such as C-phycocyanin expression or maximum growth and F_V_/F_M_ values are among other traits that can vary regardless of how closely related two strains are. In fact, strain LB1926 was challenging to test in the controlled environment conditions as it tended to form clustering even in the presence of magnetic stirring. However, using bubbling under greenhouse conditions eliminated the clustering effect.

Another interesting finding related to morphology was the linear trichomes expressed by UTEX LB 2340. This phenotype was expressed since the strain was received from UTEX at all growth conditions. We can discard stress as an explanation for the observed phenotype due to the high F_V_/F_M_ observed.

There was a correlation between values of α measured during rapid light curves and *t_1/2_* of the Kautsky curve. This strong correlation resembles those previously reported in other photosynthetic samples such as chloroplast (Briantais et al. 1979), *Chlorella* (Bates et al. 2023) and Tomato (Takayama et al. 2012) where the slow phase of the Kautsky induction curve correlated with electron transport. Although, it is well known that steady states of electron transport influence the slow phase of the Kautsky induction curve (Papageorgiou et al. 2007; Lindqvist et al. 2016), it is the first time (to our knowledge) the parameter t_1/2_ is correlated to the elements of the Platt model. The parameter t_1/2_ can potentially be used to estimate α with a higher throughput and deserves further study.

There are, nevertheless, limitations to our work. First, the use of chlorophyll *a* fluorescence method in cyanobacteria tends to be more challenging than with other photosynthetic organisms (Grigoryeva 2020; Papageorgiou et al. 2007). For example, the exchange between the PQ pool with respiratory reactions (Berry et al. 2003; Grigoryeva 2020; Papageorgiou et al. 2007), as well as the poor thylakoid stacking, yields signals that are more variable to those observed in chlorophytes and embryophytes (Sipka et al. 2021). As a consequence, determinations of F_V_/F_M_ in cyanobacteria does not directly reflect the PSII activity (like in the other photosynthetic species (Sipka et al. 2021)). However, by using the same excitation wavelength in the two types of fluorometers used, we have reduced artifacts associated with differences in optical and functional cross section as discussed by some authors (Osmond et al. 2017; Serôdio and Campbell 2021), thus, we disregard the observed result as an artifact.

Another limitation was the use of the Bradford assay, although the method is fast and sensitive (which is ideal for a phenomic study), its accuracy depends heavily on protein composition and sample purity (Compton and Jones 1985; Nasirova et al. 2026). Despite the above, the observations of this work, matched well what is known for Limnospira. The observed values of F_V_/F_M_, protein content, expression of C-phycocyanin and growth rates are well within what was reported for Limnospira grown in Zarrouk’s media (Jensen and Knutsen 1993). Thus, we disregard the introduction of any major experimental artifact that could affect the interpretation of the results presented here.

In summary, we evaluated the multiple traits of growth and photophysiology in the context of phylogenetic relations. Protein content and C-phycocyanin are parameters very conserved among species and no significant differences were found between the strains studied.

Meanwhile, we found that some traits, particularly those associated with fine tuning of photosynthesis, correlate with the phylogenetic relationships between strains, particularly within clades A and B. We discover two strains with contrasting behavior, UTEX 3086 a photosensitive strain but a high efficiency of growth at low light and temperature; while UTEX LB 2340 was light tolerant. Although some differences were able to be explained using biophysical parameters such as the magnitude of F_V_/F_M_ and velocity of fluorescence decay, these parameters were not enough to explain the observed differences in terms of growth at different environmental conditions. Other parameters such as carbon fixing, photoprotection or tolerance to photorespiration are interesting targets of future analyses when comparing the strains of *L. platensis* presented here.

## Acknowledgements

The team thanks Ing. Robert Peters for the support of this project. AZ is grateful to Profs. Tattersall and Stuart for their insightful discussions and access to infrastructure. The team is grateful for the technical support of Dr. Roepke in the isolation and quantification of phycocyanin. The team would like to thank the technical support and training of Margarita Di Profio, Irene Palumbo, Alison Smart and Dr. Aditi Das. Finally, AZ and MZ would like to thank the support of undergraduate volunteers Mateo de Lima, Abby Di Maria, Courtney Fraser and Laura Palumbo during fall 2025.

## Declarations

### Funding

AZ teams were financially supported by an NSERC Discovery Programs RGPIN-2024-04060 and DGECR-2024-00369. In addition, to the sponsorship by Track Investment Ltd, Ontario, Canada and Brock University internal funds as part of the start-up package of Dr. Zavafer.

### Competing interests

The authors declare that they have no known competing financial interests or personal relationships that could have appeared to influence the work reported in this paper.

### Availability of data and material

Data available at Federated Research Data Repository https://www.frdr-dfdr.ca

## Author’s contributions

### CRediT statement

Conceptualization: AZ; Methodology: AZ; Software: AZ; Validation: AZ, MZ; Formal analysis: AZ, MZ; Investigation: MZ, AZ; Resources: APL, AZ; Data Curation: CP, AZ, MP; Writing - Original Draft: MZ, AZ; Writing - Review & Editing: AZ, CP, APL ; Visualization: AZ; Supervision: AZ; Project administration: AZ; Funding acquisition: CP, AZ.

### Corresponding author

Alonso Zavafer

## Notes

### Competing Interest Statement

The authors have declared no competing interest.

