## Supplementary material for "COMPARATIVE PHENOMICS IN *Limnospira platensis* STRAINS REVEALS PHOTOBIOENERGETIC SIMILARITIES AND DIFFERENCES IN AMERICAN AND AFRICAN STRAINS": Supp data

SUPPLEMENTARY DATA

Mateo Zavala [1,3], Jose Armando Perez-Loera [1,2], Carli Peters [1,5], A Zavafer [1,2,3]

[1] Department of Biological Sciences, Brock University, St. Catharines, Ontario, Canada

[2] Department of Physics, Brock University, St. Catharines, Ontario, Canada

[3] Brock University's Microphyte Collection, Brock University, St. Catharines, Ontario, Canada

[4] Yousef Haj-Ahmad Dept. of Engineering, Brock University, St. Catharines, Ontario, Canada

[5] Big Algae Ltd, St. Catharines, Ontario, Canada

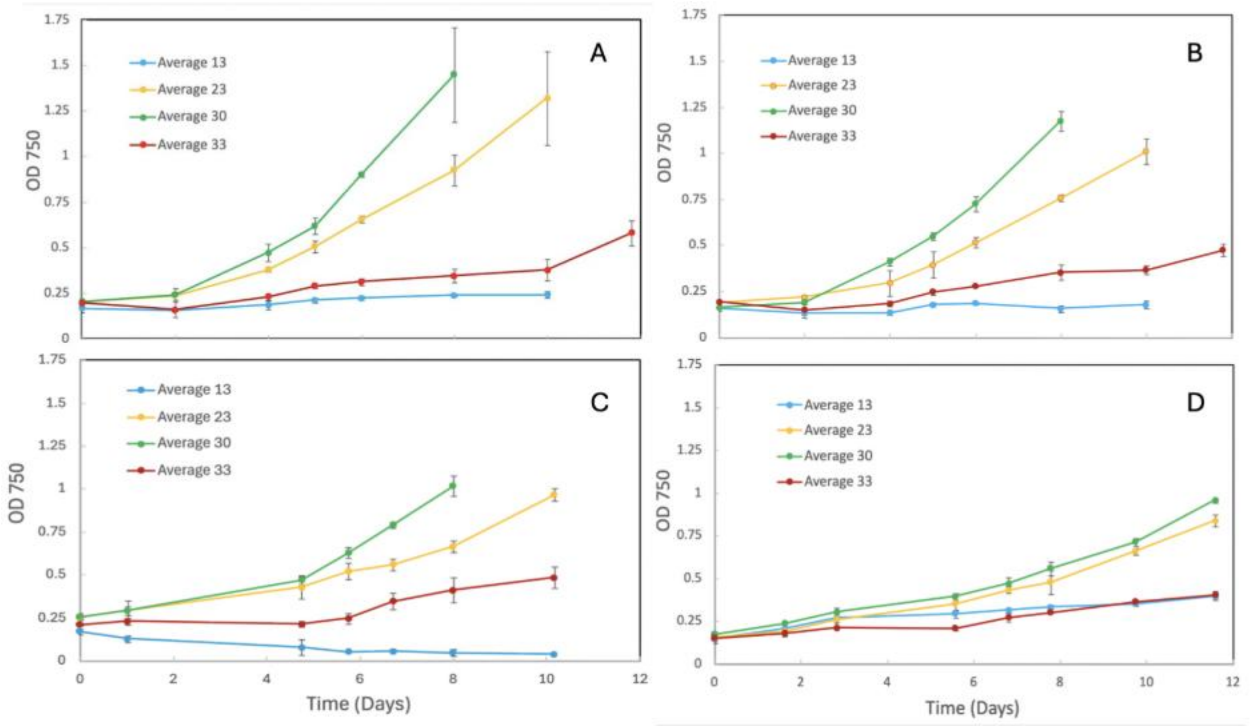


**Figure S1** Optical density ($OD_{750}$) measurements over 13 days showing the growth of spirulina strains in different temperatures. (A) Strain 3086, (B) strain LB2342, (C) strain LB2340, and (D) strain LB1928. Each panel compares growth across the indicated temperature conditions. *n*=4

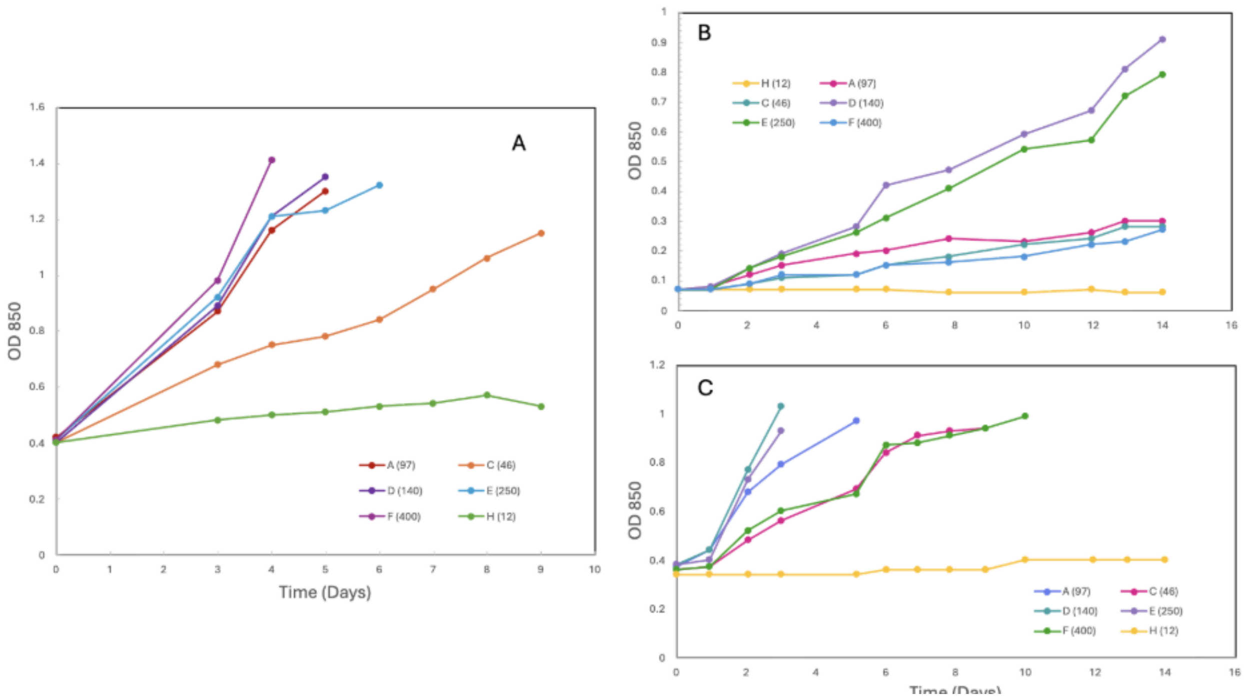


**Figure S2** Growth curves ($OD_{850}$ vs time (days)) for the determination of optimal light intensity in strain 3086 across two experimental rounds. Round 1 is shown on the left and round 2 on the right. (A) Growth curve for round 1, measured directly in the culture using a Roux bottle T25 (maximum volume 40 mL). (B) Growth curve for round 2, measured from 1 mL aliquots using a cuvette with an optical path length of 5 mm. (C) Growth curve for round 2, measured directly in the culture using a Roux bottle T25 (maximum volume 40 mL).

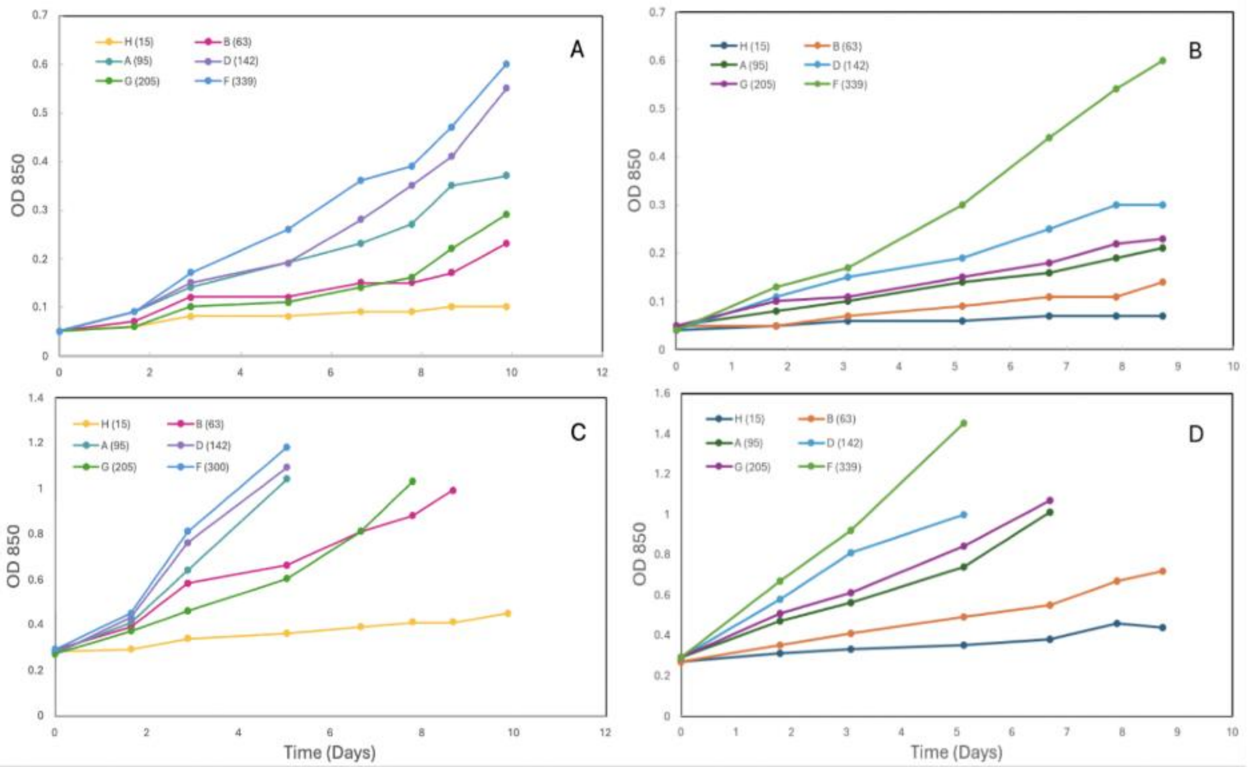


**Figure S3** Growth curves ($OD_{850}$ vs time (days)) for the determination of optimal light intensity in strain LB2342 across two experimental rounds. Round 1 is shown on the left and round 2 on the right. (A) Growth curve for round 1, measured from 1 mL aliquots using a cuvette with an optical path length of 5 mm. (B) Growth curve for round 2, measured from 1 mL aliquots using a cuvette with an optical path length of 5 mm. (C) Growth curve for round 1, measured directly in the culture using a Roux bottle T25 (maximum volume 40 mL). (D) Growth curve for round 2, measured directly in the culture using a Roux bottle T25 (maximum volume 40 mL).

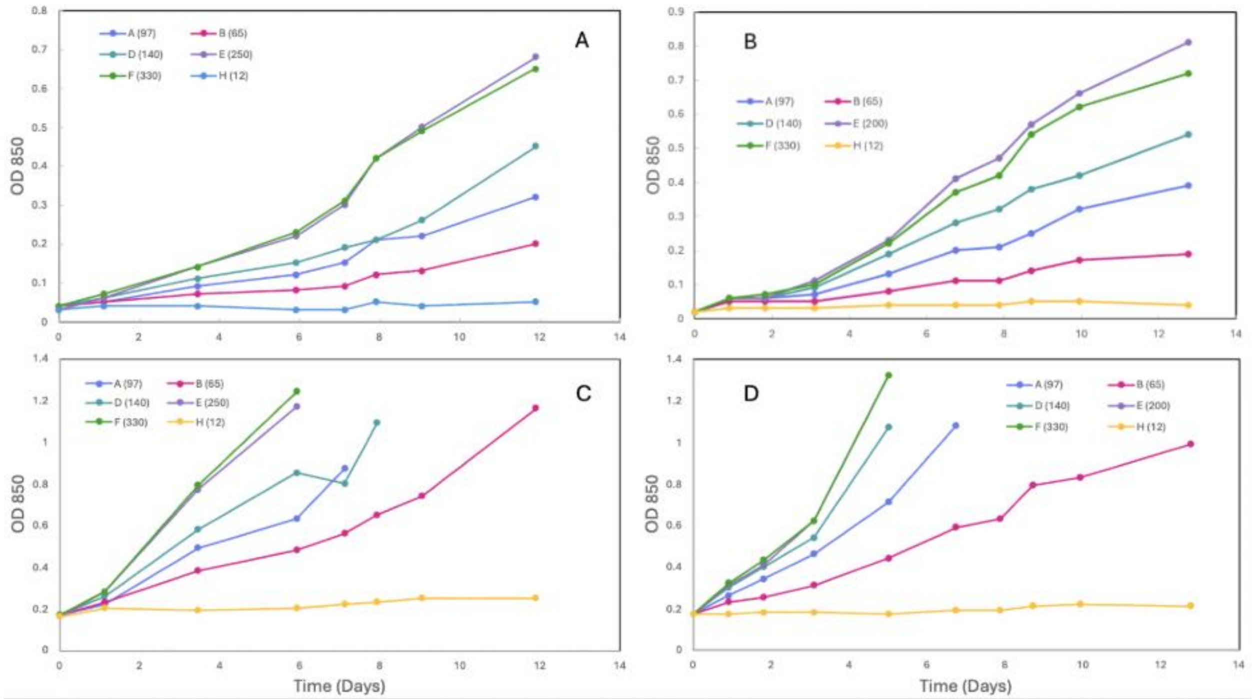


**Figure S4** Growth curves ($OD_{850}$ vs time (days)) for the determination of optimal light intensity in strain LB2340 across two experimental rounds. Round 1 is shown on the left and round 2 on the right. (A) Growth curve for round 1, measured from 1 mL aliquots using a cuvette with an optical path length of 5 mm. (B) Growth curve for round 2, measured from 1 mL aliquots using a cuvette with an optical path length of 5 mm. (C) Growth curve for round 1, measured directly in the culture using a Roux bottle T25 (maximum volume 40 mL). (D) Growth curve for round 2, measured directly in the culture using a Roux bottle T25 (maximum volume 40 mL).

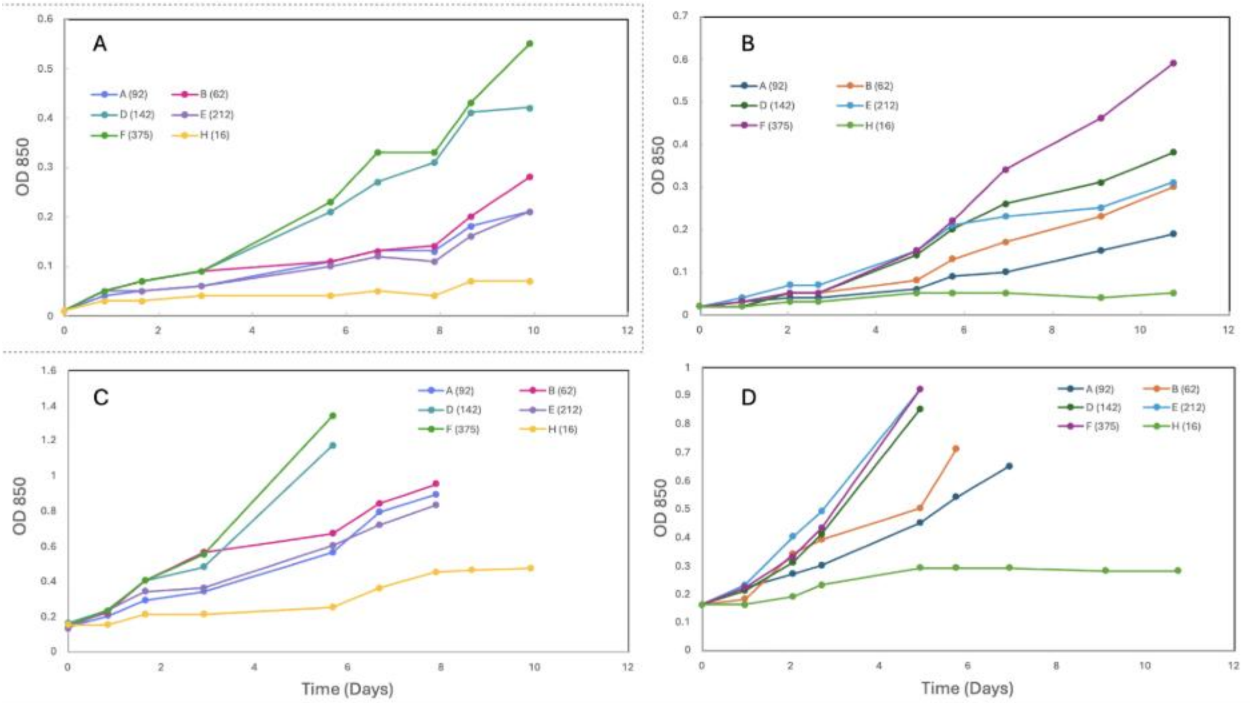


**Figure S5** Growth curves ($OD_{850}$ vs time (days)) for the determination of optimal light intensity in strain LB1928 across two experimental rounds. Round 1 is shown on the left and round 2 on the right. (A) Growth curve for round 1, measured from 1 mL aliquots using a cuvette with an optical path length of 5 mm. (B) Growth curve for round 2, measured from 1 mL aliquots using a cuvette with an optical path length of 5 mm. (C) Growth curve for round 1, measured directly in the culture using a Roux bottle T25 (maximum volume 40 mL). (D) Growth curve for round 2, measured directly in the culture using a Roux bottle T25 (maximum volume 40 mL).

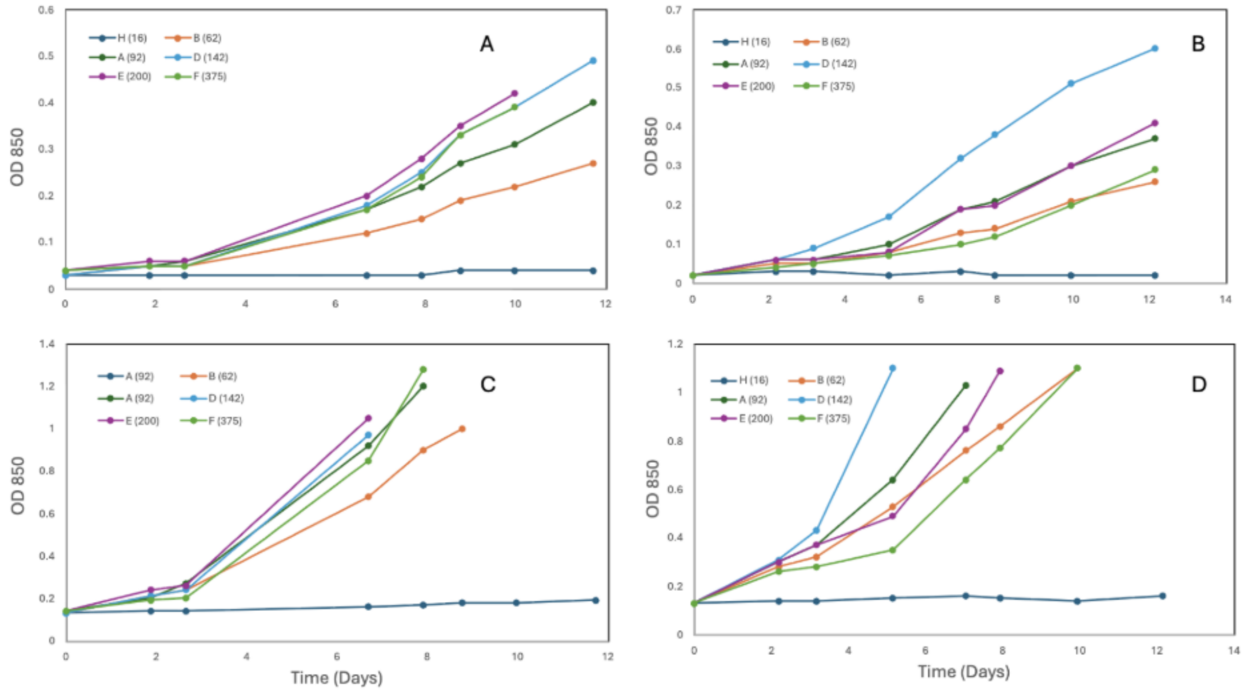


**Figure S6** Growth curves ($OD_{850}$ vs time (days)) for the determination of optimal light intensity in strain LB1926 across two experimental rounds. Round 1 is shown on the left and round 2 on the right. (A) Growth curve for round 1, measured from 1 mL aliquots using a cuvette with an optical path length of 5 mm. (B) Growth curve for round 2, measured from 1 mL aliquots using a cuvette with an optical path length of 5 mm. (C) Growth curve for round 1, measured directly in the culture using a Roux bottle T25 (maximum volume 40 mL). (D) Growth curve for round 2, measured directly in the culture using a Roux bottle T25 (maximum volume 40 mL).

**Table S1** Calibration equations relating optical density (OD) to biomass for spirulina strains cultivated under greenhouse conditions. OD measurements were obtained at different wavelengths and optical path lengths. For each condition, the productivity after 25 days, the slope, intercept, and coefficient of determination ($R^2$) of the linear fitted equations are reported.

| Strain | Productivity | λ | **Optical path** | **Slope** | **Intercept** | $R^2$ |
|---|---|---|---|---|---|---|
| Units | g/L | nm | mm | L/g | | |
| 3086 | 1.883 | 750 | 10 | 1.104 | 0.159 | 0.919 |
| | | | 5 | 0.586 | 0.039 | 0.963 |
| | | 850 | 10 | 1.040 | 0.115 | 0.933 |
| | | | 5 | 0.510 | 0.034 | 0.964 |
| LB 2342 | 1.903 | 750 | 10 | 1.104 | 0.159 | 0.919 |
| | | | 5 | 0.586 | 0.039 | 0.963 |
| | | 850 | 10 | 1.040 | 0.115 | 0.933 |
| | | | 5 | 0.510 | 0.034 | 0.964 |
| LB 2340 | 1.889 | 750 | 10 | 0.942 | 0.328 | 0.915 |
| | | | 5 | 0.518 | 0.090 | 0.932 |
| | | 850 | 10 | 0.942 | 0.188 | 0.952 |
| | | | 5 | 0.493 | 0.067 | 0.972 |
| LB 1928 | 1.711 | 750 | 10 | 1.099 | 0.202 | 0.983 |
| | | | 5 | 0.587 | 0.041 | 0.978 |
| | | 850 | 10 | 1.014 | 0.156 | 0.984 |
| | | | 5 | 0.515 | 0.037 | 0.973 |
| LB 1926 | 1.585 | 750 | 10 | 1.313 | 0.029 | 0.974 |
| | | | 5 | 0.677 | 0.010 | 0.971 |
| | | 850 | 10 | 1.232 | 0.013 | 0.973 |
| | | | 5 | 0.595 | 0.004 | 0.975 |

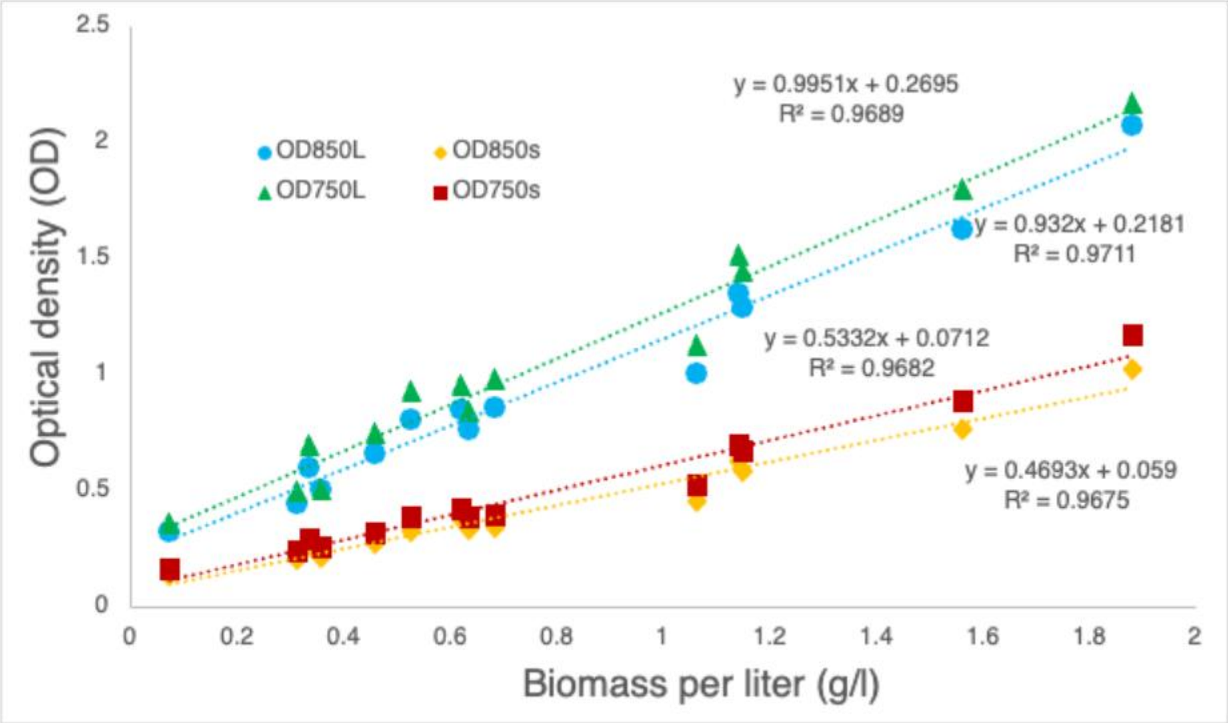


**Figure S7** Calibration curves of $OD_{750}$ and $OD_{850}$ versus dry biomass (g/L) for strain 3086 at optical path lengths of 5 mm (s) and 10 mm (L), with the corresponding linear regression equations and coefficients of determination ($R^2$) for each condition.

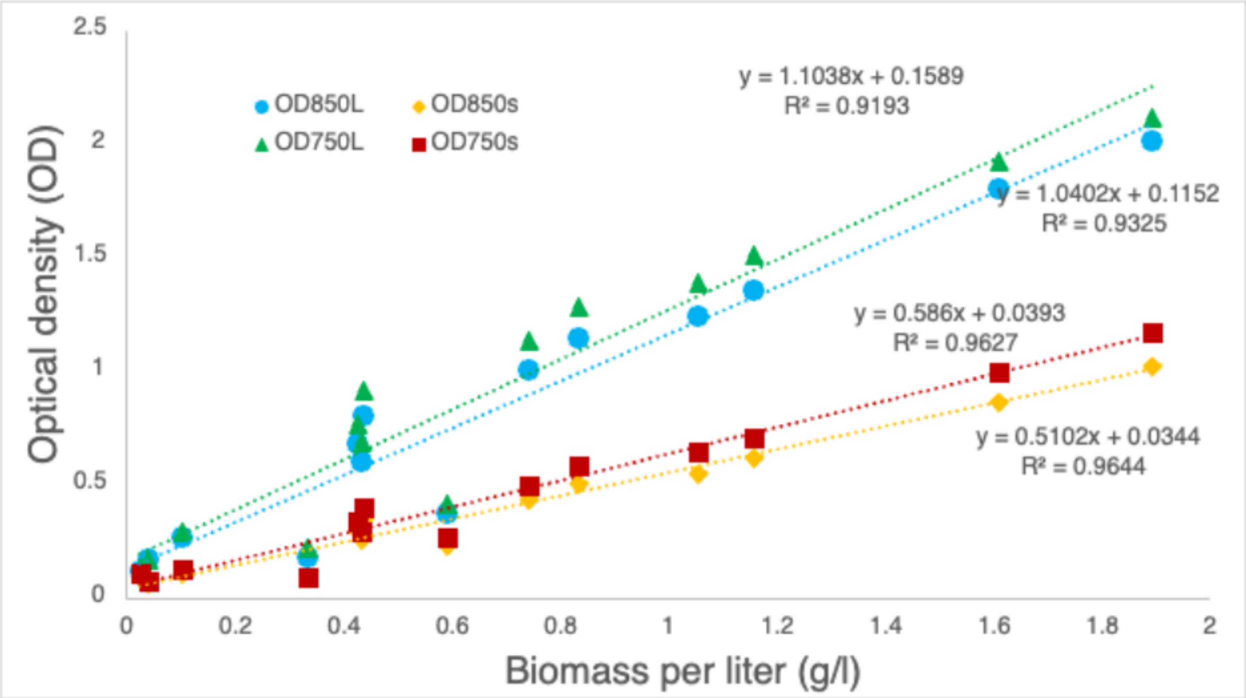


**Figure S8** Calibration curves of $OD_{750}$ and $OD_{850}$ versus dry biomass (g/L) for strain LB2342 at optical path lengths of 5 mm (s) and 10 mm (L), with the corresponding linear regression equations and coefficients of determination ($R^2$) for each condition.

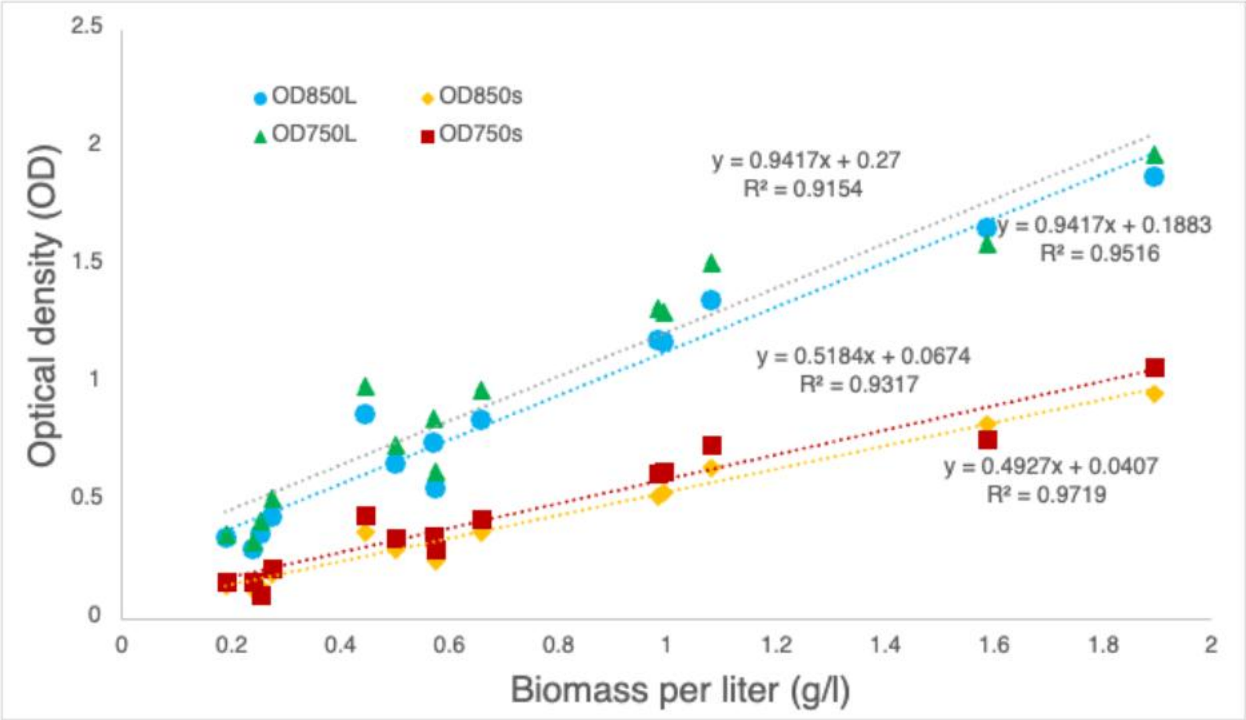


**Figure S9** Calibration curves of $OD_{750}$ and $OD_{850}$ versus dry biomass (g/L) for strain LB2340 at optical path lengths of 5 mm (s) and 10 mm (L), with the corresponding linear regression equations and coefficients of determination ($R^2$) for each condition.

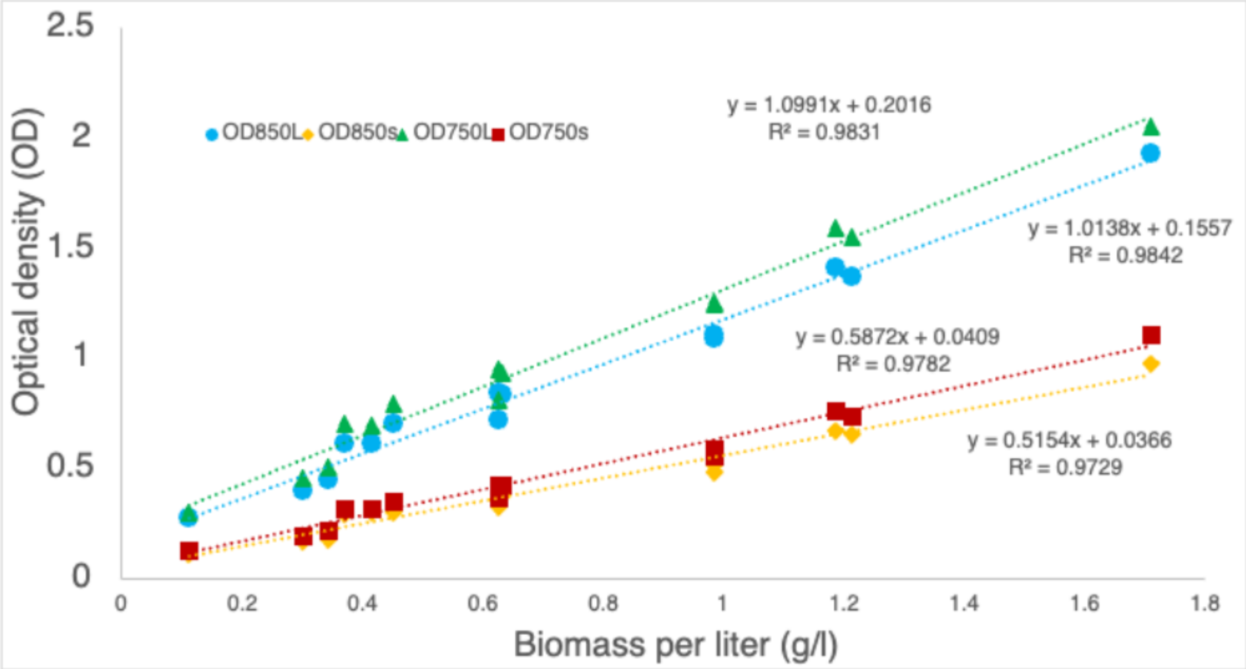


**Figure S10** Calibration curves of $OD_{750}$ and $OD_{850}$ versus dry biomass (g/L) for strain LB1928 at optical path lengths of 5 mm (s) and 10 mm (L), with the corresponding linear regression equations and coefficients of determination ($R^2$) for each condition.

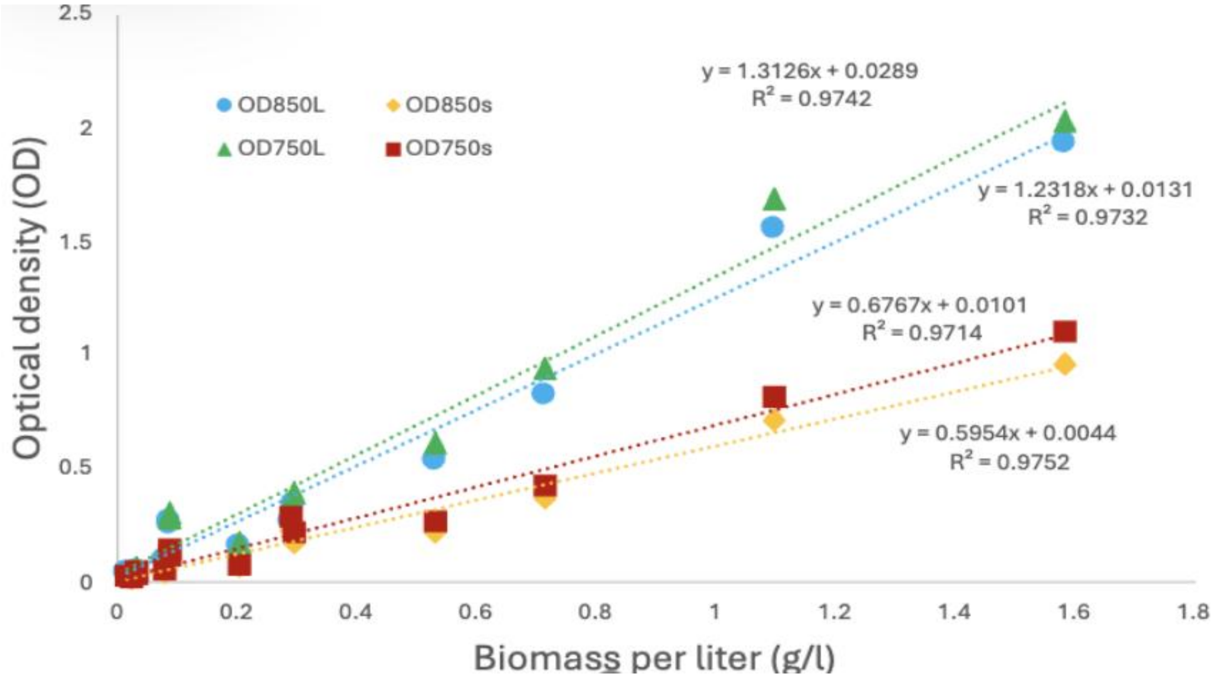


**Figure S11:** Calibration curves of $OD_{750}$ and $OD_{850}$ versus dry biomass (g/L) for strain LB1926 at optical path lengths of 5 mm (s) and 10 mm (L), with the corresponding linear regression equations and coefficients of determination ($R^2$) for each condition
